# Elucidating the Transport Mechanisms and Metabolic Roles of Serine, Threonine, and Glycine in *Trypanosoma cruzi*

**DOI:** 10.1101/2024.06.29.601350

**Authors:** Mayke Bezerra Alencar, Richard Marcel Bruno Moreira Girard, Marcell Crispim, Carlos Gustavo Baptista, Marc Biran, Frederic Bringaud, Ariel Mariano Silber

## Abstract

l-Serine (l-Ser) and l-Threonine (l-Thr) have versatile roles in metabolism. In addition to their use in protein synthesis, these amino acids participate in the biosynthesis pathways of other amino acids and even phospholipids. Furthermore, l-Ser and l-Thr can be substrates for a Ser/Thr dehydratase (Ser/ThrDH), resulting in pyruvate (Pyr) and 2-oxobutyrate, respectively, thus being amino acids with anaplerotic potential. *Trypanosoma cruzi*, the etiological agent of Chagas disease, uses amino acids in several biological processes: metacyclogenesis, infection, resistance to nutritional and oxidative stress, osmotic control, etc. In this study, we investigated the import and metabolism of l-Ser, l-Thr, and Gly in *T. cruzi*. Our results demonstrate that these amino acids are transported from the extracellular environment into *T. cruzi* cells through a saturable transport system that fits the Michaelis-Menten model. Our results show that l-Ser and l-Thr can sustain epimastigote (Epi) cell viability under nutritional stress (NS) conditions and can stimulate oxygen consumption to maintain intracellular ATP levels. Additionally, our findings indicate that l-Ser plays a role in establishing the mitochondrial membrane potential (ΔΨ_m_) in *T. cruzi*. l-Ser is also involved in energy metabolism via the Ser-Pyr pathway, which stimulates the production and subsequent excretion of acetate and alanine. Our results demonstrate the importance of l-Ser and l-Thr in the energy metabolism of *T. cruzi* and provide new insights into the metabolic adaptations of this parasite during its life cycle.

## INTRODUCTION

*Trypanosoma cruzi* is the etiological agent of Chagas disease, a.k.a American Trypanosomiasis. During its life cycle, the parasite transitions between different environments of different hosts, such as the mammalian host-cell cytoplasm, the mammalian blood and different regions of the reduviid insect’s digestive tube (midgut and rectum lumen). To survive in these environments, *T. cruzi* reprograms its metabolism, being able to metabolize various carbon/energy sources (1–5). These parasités habitats range from those with abundant to those with low carbohydrates availability, such as the intestinal tract of the starved insect vector (6–10). In this regard, it has been well described that these parasites can switch between the consumption of carbohydrates and that of amino acids and fatty acids (11–14). To cope with the scarcity of metabolites in these different environments *T. cruzi* must acquire and utilize the available nutrients as carbon and energy sources [reviewed in: (15)].

Among the available nutrients in most environments colonized by *T. cruzi* throughout its life cycle, l-Ser, l-Thr, and Gly are almost omnipresent. Despite this, few groups have devoted themselves to studying in depth the metabolism of these amino acids in *T. cruzi*. In the 1970s, Hampton determined the participation of l-Ser in the production of CO_2_, indicating that this molecule is metabolizable by the parasite (16). Additionally, there is already extensive evidence of the participation of l-Thr in the mitochondrial metabolism of *Trypanosoma brucei* (17–19). Although there is no evidence of the participation of Gly in energy metabolism, the carbons of Gly may contribute to the formation of l-Ser through the reversible activity of serine hydroxymethyltransferase (SHMT) (20,21). Therefore, Gly could be metabolized via SHMT, thus participating in mitochondrial ATP production.

Despite the demonstrated potential of these amino acids to trigger ATP biosynthesis in many organisms (protists, bacteria, yeast, and mammals) (17,19,22–30), their consumption and metabolism in *T. cruzi* remain almost unexplored. In this work, we explore the uptake of l-Ser, l-Thr, Gly, and their role in the parasités bioenergetics and nutritional stress (NS) resilience.

## RESULTS

### *T. cruzi* epimastigote transport l-Ser, l-Thr and Gly from the extracellular medium

To characterize the l-Ser, l-Thr and Gly uptake, their transport was measured as a function of time. For this, the parasites were incubated with each radioactively traced amino acid at a 5 mM concentration, assuming that this concentration is saturating for each transport system. The transport of these metabolites along time was fittable to an exponential decay function (l-Ser: *r*^2^ = 0.99, l-Thr: *r*^2^ = 0.98 and Gly: *r*^2^ = 0.98), as expected for saturable protein-mediated transport systems (Fig. 1A, B and C). At initial times, the resulting exponential-decay functions can be assumed as near-linear, allowing a simple and reliable calculation of initial velocities for substrate uptake. Our data show that linear functions can be accurately fitted to the measured uptake of l-Ser, l-Thr, and Gly until 10 min (*r*^2^= 0.96, *r*^2^= 0.95, and *r*^2^= 0.94 for l-Ser, l-Thr, and Gly, respectively (insets in Fig. 1 A, B and C)). Therefore, 3 min was set as the time window to measure the initial velocity (*V*_0_) of l-Ser, l-Thr, and Gly uptake.

**Figure 1.**
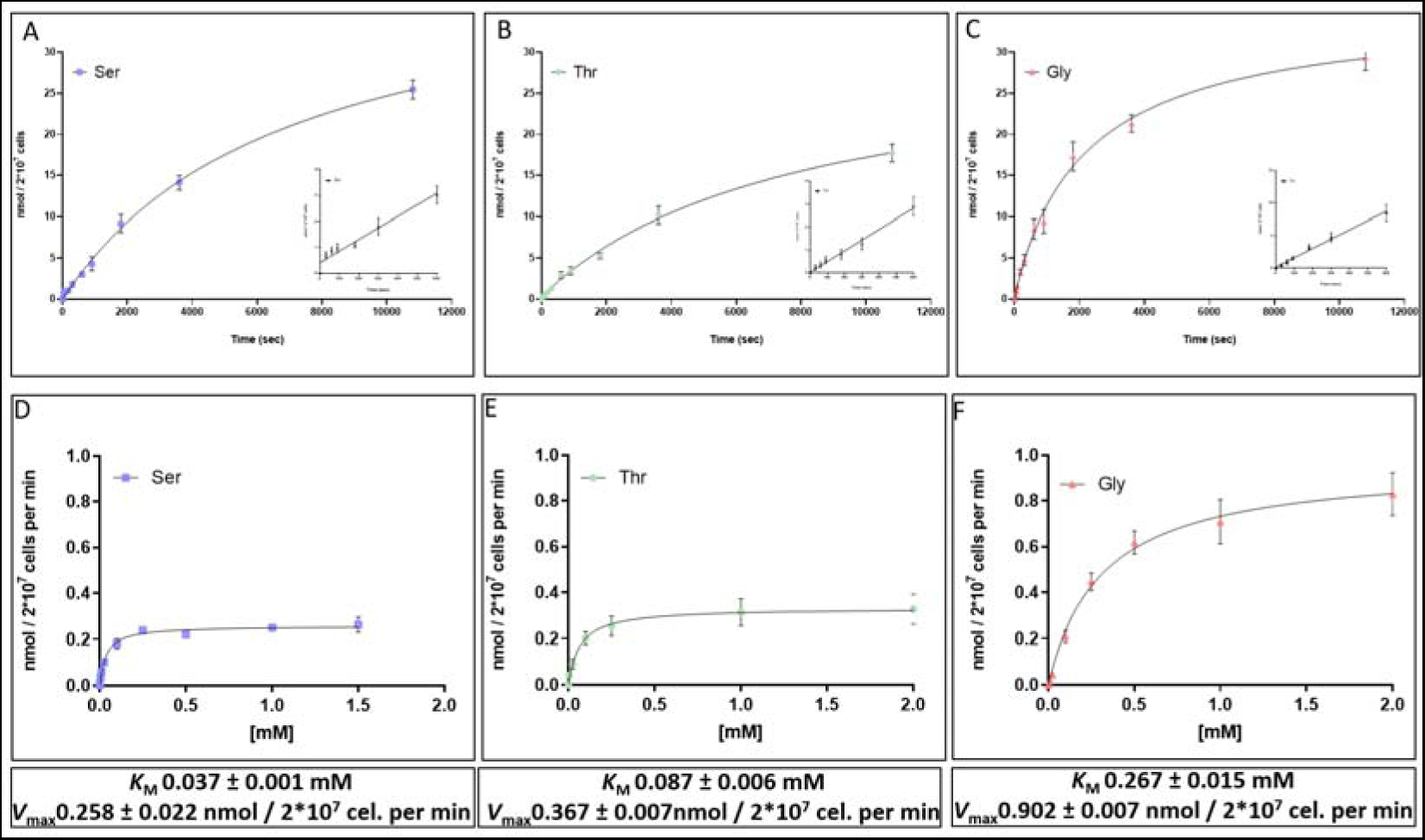
Uptake of l-Ser, l-Thr, and Gly in *T. cruzi* epimastigote as a function of time and concentration. (A, B, and C) Time-course transport of 5 mM l-Ser, l-Thr, and Gly, respectively. (D, E, and F) Uptake of l-Ser, l-Thr, and Gly, respectively, as a function of their concentration. The insets represent the adjustment of incorporation as a function of time to a linear function. The table below the graphs gives the values of the kinetic parameters calculated by the Michaelis-Menten model. *r*^2^ values of 0.96, 0.95, and 0.94 for l-Ser, l-Thr, and Gly, respectively.

To analyze the kinetics for the transport of each amino acid, epimastigote forms (Epi) of the parasites were incubated with different concentrations of l-Ser, l-Thr or Gly for 3 min and the uptake of each amino acid was measured (through the cell-associated radioactivity). The hyperbolic functions corresponding to the classic Michaelis-Menten kinetics fit the data corresponding to each amino acid. The non-linear hyperbolic functions that best explained our data showed robust adjustments (*r*^2^ = 0.96, 0.95, and 0.94 for l-Ser, l-Thr, and Gly, respectively). The values for *K*_M_ and *V*_max_ were derived from these functions for the three substrates (Fig. 1 D, E, and F). Summarizing, the results showed that l-Ser, l-Thr, and Gly are taken up from the extracellular medium by *T. cruzi* Epi through saturable transport systems and their kinetics fit the Michaelian model. Therefore, the *K*_M_ and *V*_max_ were calculated as being 37 ± 0.001 µM, 87 ± 0.006 µM,267 ± 0.015 µM and 0.258 ± 0.022 nmol min^-1^ 2 x10^7^ cells, 0.367 ± 0.007 nmol min^-1^ 2 x10^7^ cells and 0.902 ± 0.007 nmol min^-1^ 2 x10^7^ cells for l-Ser, l-Thr, and Gly, respectively (Fig. 1 D, E, and F).

Then, the possible ability of l-Ser, l-Thr, and Gly to share the same transport system was investigated. To this end, a cross-competition assay was performed to assess the inhibition rate of each amino acid transport in the presence of the potentially competing amino acid at a concentration corresponding to 10-fold the *K*_M_ value (370 µM, 870 µM and 2600 µM for l-Ser, l-Thr and Gly, respectively) (Table 1). The uptake of l-Ser was diminished by 50% and 45%, respectively, by l-Thr and Gly, while the uptake of l-Thr was diminished by 82% and 42%, respectively, by l-Ser and Gly, and the transport of Gly was inhibited by 83% and 58%, respectively, by l-Ser and l-Thr. Our findings show that l-Ser, l-Thr, and Gly indeed compete with each other, and thus, support the hypothesis of a shared transporter for these three amino acids. l-Ser uptake was set as a proxy for further characterizing the activity for all three. Using the same conditions, the ability of other amino acids to impair l-Ser transport was assessed. While structurally related amino acids such as d-Ser, l-Cys, d-Ala and l-Ala resulted in partial inhibition of l-Ser transport, structurally dissimilar amino acids, such as branched-chain amino acids (BCAA; l-Leu, l-Ile and l-Val) (31), l-His (32) and l-Phe did not significantly affect it, as expected (Table 1).

**Table 1.**
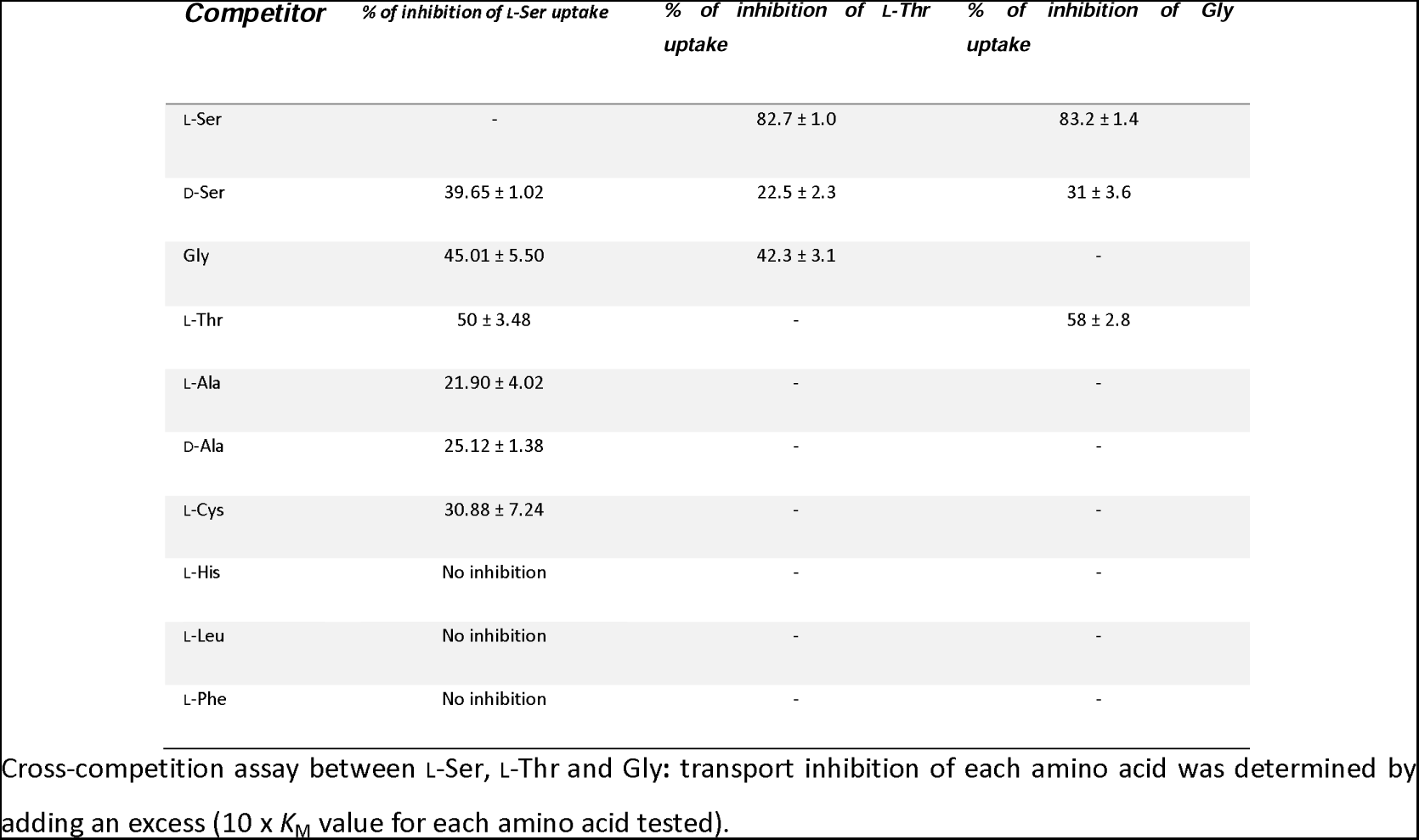
Percentage of inhibition in transport observed in cross-competition assay.

### Determination of the driving force for l-Ser uptake

To investigate the effect of intracellular ATP levels on this transport system, l-Ser uptake was measured in parasites that were treated for 30 min with oligomycin A (Oly), a known F_O_-ATP synthase inhibitor which produces intracellular ATP depletion (31), or without (control). A control group treated with Oly immediately before measuring the l-Ser uptake was included, to discard an off-target effect of Oly on the l-Ser transport. Cells pre-incubated with Oly for 30 min showed a l-Ser uptake decreased by approximately 40.3 ± 13%, while l-Ser uptake levels were unaffected without pre-incubation when compared with the untreated control. To determine whether the transport system was dependent on the proton gradient (Δp) across the plasma membrane, the *V*_0_ of l-Ser incorporation in the presence of the protonophore CCCP was measured. CCCP has two known effects on cells: i) it collapses the H^+^-related membrane potential (Δ_Ψ_); and ii) it depletes intracellular ATP levels by reversing the mitochondrial F_1_F_o_-ATP synthase reaction. To distinguish these two effects on the transport assay, the l-Ser transport was measured also in the presence of CCCP supplemented with Oly (to avoid ATP hydrolysis by reversal of the F_1_F_o_-ATP synthase activity). CCCP, both, in the presence and absence of Oly, significantly diminished the transport activity when compared to the positive control (C+), indicating that this transport system is dependent on the Δp across the plasma membrane, which in turn depends on the intracellular ATP levels (Fig. 2 and Table 2).

**Figure 2.**
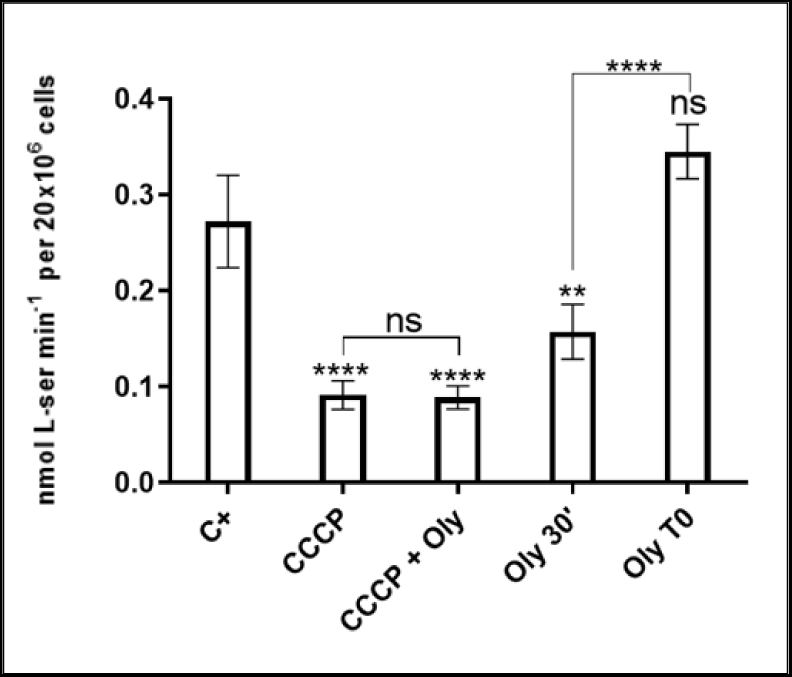
(A) Effect of oligomycin A and CCCP on l-Ser uptake: the dependence of l-Ser transport on intracellular ATP levels (Oly 30’) and the H^+^ gradient (CCCP) was assessed. CCCP can rapidly trigger ATP hydrolysis by mitochondrial ATPase to reestablish the H^+^ gradient, leading to ATP depletion in the cell. To distinguish between these phenomena, the parasites were incubated with CCCP in the presence and absence of Oly (CCCP + Oly). To control for non-specific inhibition of the transport system by Oly, we added it without pre-incubation (Oly T0). These graphs represent three independent biological replicates. Statistical analysis was performed using one-way ANOVA with Tukey’s post-test (**** P< 0.0001; ** P: 0.002).

**Table 2.** Inhibition of l-Ser uptake in the presence of FCCP and Oly.

| Treatment | % of L-Ser uptake inhibition |
| --- | --- |
| 25 μM CCCP | 63.8 ± 9 |
| 25 μM CCCP + 0.5 μg/μL Oly | 66.3 ± 0.2 |
| 0.5 μg/μL Oly 30' | 40.3 ± 13 |
| 0.5 μg/μL Oly T0 | nd* |

\*not detected.

### Thermodynamic analysis of l-Ser uptake

To assess the effect of temperature on l-Ser uptake, the *V*_max_ was measured at different temperatures ranging between 4 °C and 60 °C, which showed a maximum at 40 °C. In addition, under these conditions the changes in *V*_max_ in the exponential region of the curve were utilized to calculate a Q_10_ of 1.948, indicating a temperature-dependent process. Additionally, *V*_0_ measurements made between 4 °C and 60 °C (R^2^: 0.988) were used to calculate an activation energy (*E*_a_) of 51.2 ± 2.4 kJ/mol (Fig. 3 A and B). Additionally, from the Arrhenius equation, it was possible to calculate the transporter turnover, which allowed an estimation of the number of transporters being used for the substrate uptake as 0.099 attomol per cell (see Text S1 in the supplemental material). This is equivalent to approximately 6 x 10^4^ active sites per cell.

**Figure 3.**
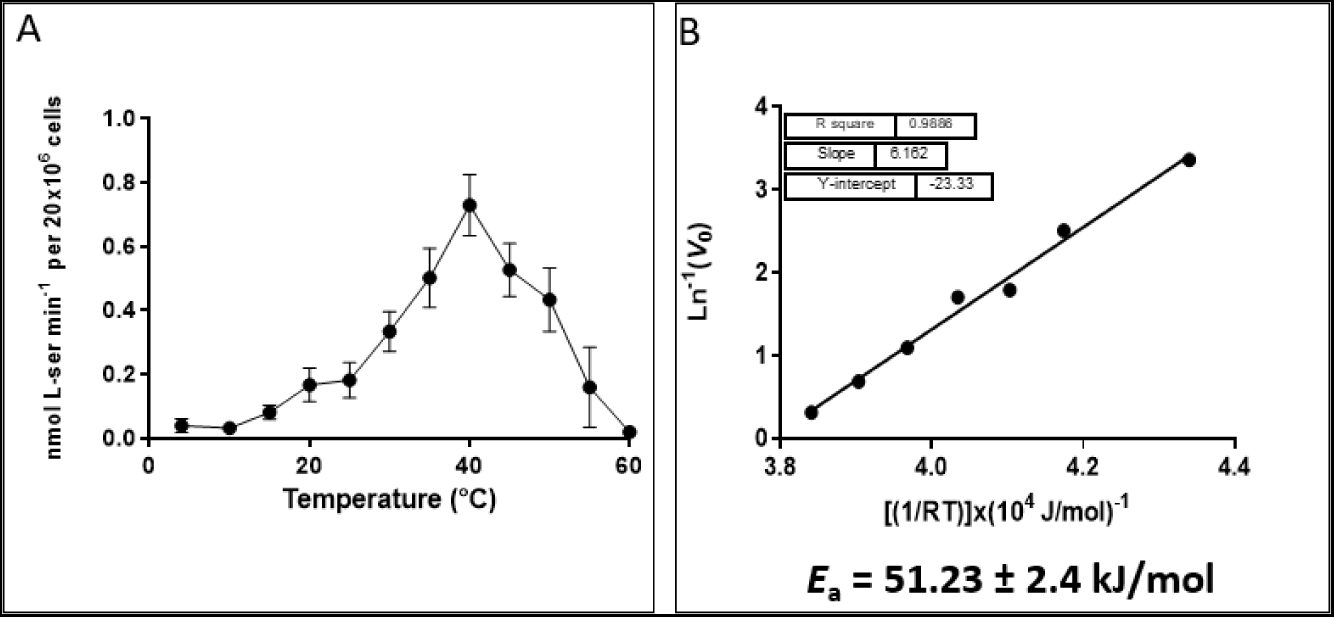
Effect of temperature on l-Ser uptake: (A) The *V*_0_ was measured at saturating concentrations of l-Ser after 3 min of uptake at specific temperatures. (B) An Arrhenius plot was created by performing a linear fit between the *V*_0_ values measured at temperatures ranging from 4 to 40 °C. These graphs represent three independent biological replicates.

### Uptake of l-Ser in the different forms of the *T. cruzi* life cycle

Previous research on changes in nutrient uptake profiles during *T. cruzi* life cycle has been scarce. However, two works have previously shown that the uptake of glucose (Glc) (33) and l-glutamine (34) change according to the life-cycle stage of the parasite, indicating metabolic adaptations to different environments. Herein, the uptake of l-Ser in vertebrate intracellular stages (Amastigote - Ama and Intracellular Epimastigote - Epi-like), stages present in the insect vector (Epi - Epimastigote and Metacyclic Trypomastigote - MTrypo), and those present in the bloodstream of vertebrates (Bloodstream Trypomastigote - BTrypo) was measured. The obtained results show that l-Ser transport varies among the parasite’s life-cycle stages. Interestingly, the intracellular Epi-like stage captured approximately 5 times more l-Ser from the medium than Epi forms, while MTrypo and BTrypo capture 5 times less than Epi (Fig. 4).

**Figure 4.**
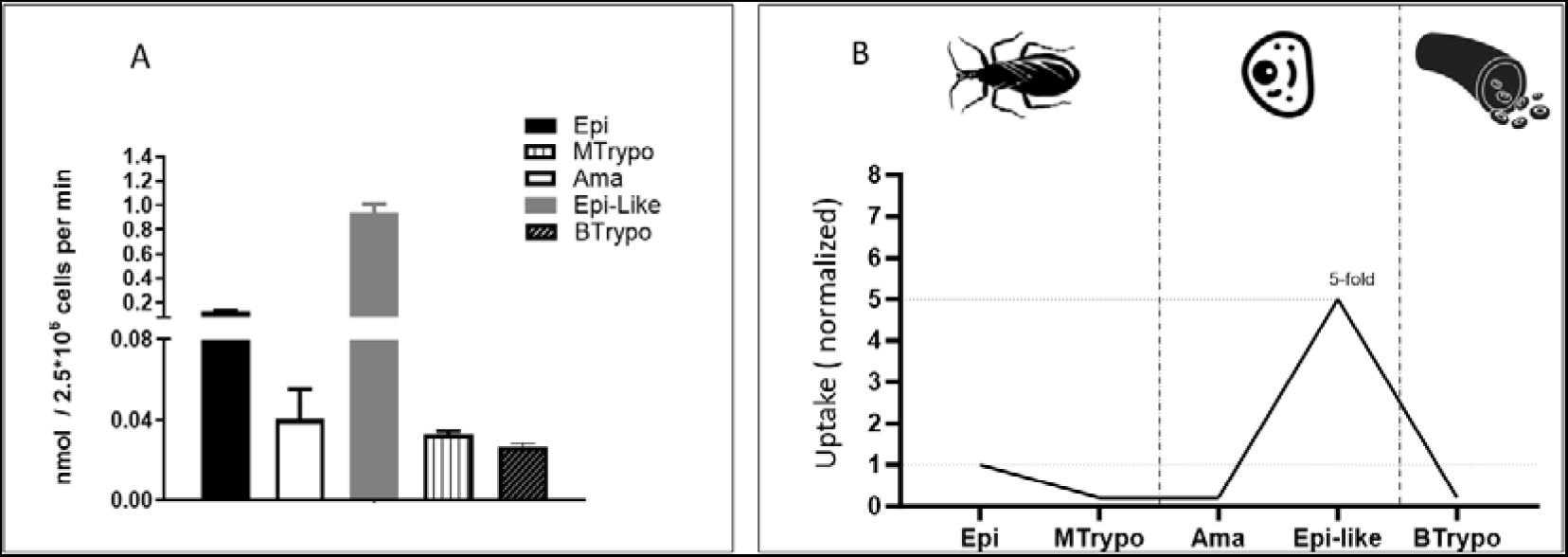
Transport of l-Ser in different life-cycle stages of *T. cruzi*. A) Transport of l-Ser; B) Transport of l-Ser (normalized). Epi: Epimastigote; Ama: Amastigote; Epi-like: Epimastigote-like; Meta: Metacyclic; Trypo: Bloodstream trypomastigote. The data are representative of three independent biological triplicates. The panel on the right represents the normalized transport rates, considering the transport measured in epimastigotes as 100%.

### Bioenergetics of l-Ser, l-Thr and Gly in *T. cruzi* epimastigotes

To investigate the bioenergetic fate of the analyzed amino acids, and their potential role in cell survival during NS was examined. Cells were incubated in PBS supplemented or not (as negative control) with 5 mM of the amino acids under study. Cells supplemented with l-His (32) or l-His plus Glc were used as positive controls. After 48h of NS, only l-Thr succeeded in maintaining cell viability at the level of the positive control (His). However, during longer NS periods (72h and 96h), Gly and l-Ser can sustain viability significantly better than only PBS. Taken together, these results indicate that all three metabolites can protect the cells from NS, with l-Thr being more efficient than l-Ser and Gly (Fig. 5 A).

**Figure 5.**
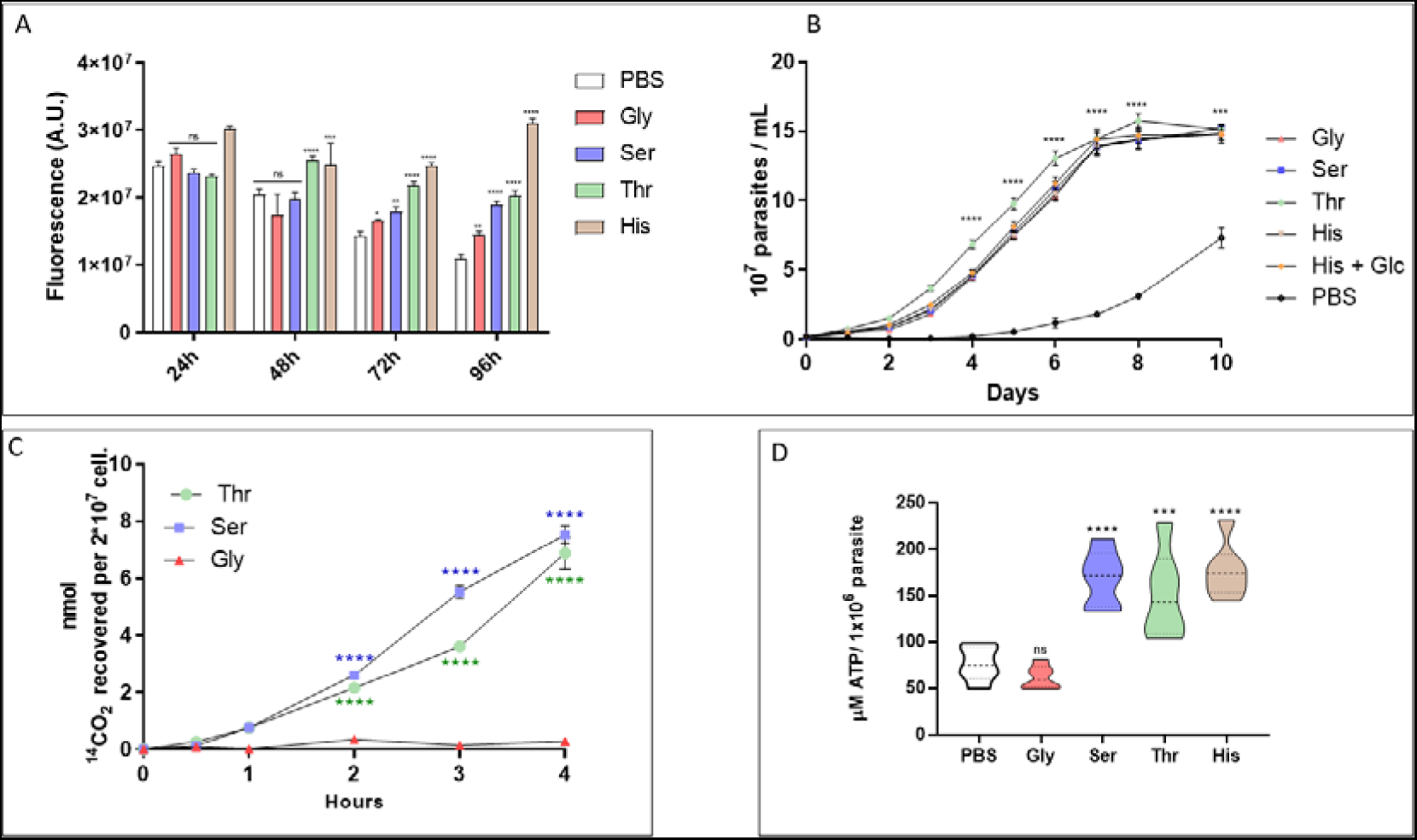
(A) Viability assay of *T. cruzi* epimastigotes as a function of nutritional stress: viability was measured by the irreversible reduction of resazurin to resorufin. His (5 mM) was used as a positive control, and no exogenous source of carbon was used as a negative control. (B) Recovery assay of *T. cruzi* epimastigotes under NS conditions: the proliferation profile was evaluated after 72 h of NS in the presence or absence (only PBS) of exogenous carbon sources, 5 mM His and l-His + Glc (5 mM each, positive controls) and 5 mM Gly, l-Ser and l-Thr. Calibration curves were performed using known parasite densities. As negative control, PBS without exogenous carbon sources was used. (C) Production of ^14^CO_2_ by breaking down Gly, l-Ser and l-Thr in *T. cruzi* epimastigote. (D) Intracellular levels of ATP in epimastigotes recovered from NS with Gly, l-Ser and l-Thr: after 16 h of NS, the parasites were recovered for 3 h in the presence (or not, negative control – only PBS) of 5 mM His (positive control) Gly, l-Ser and or l-Thr. ATP levels were measured by detecting luminescence in a coupled luciferase reaction. The graphs are representative of three independent biological replicates. Two-way ANOVA with Tukey’s post-test was used for statistical analysis: ** P<0.0029; *** P<0.0007; **** P<0.0001.

The fact that l-Ser, l-Thr and Gly were able to keep the cells alive under NS conditions does not necessarily imply their capacity to resume proliferation when re-incubated in rich LIT medium. Thus, this possibility was investigated by performing a proliferation recovery experiment following NS. Our results showed that l-Ser, l-Thr, and Gly can maintain the proliferative capacity of Epi forms after arrest induced by NS, at similar levels as the positive control. These results strongly indicate that these amino acids can assist the parasite to cope with NS (Fig. 5 B).

In most organisms, l-Ser and l-Thr carbons primarily contribute to energy metabolism via the Ser-pyruvate (Pyr) and Thr-Acetyl-CoA pathways respectively. Once these metabolites enter the tricarboxylic acid (TCA) cycle, they can be broken down into CO_2_. To investigate this possibility, Epi forms were incubated with l-[3-^14^C]Ser, l-[U-^14^C]Thr, and [U-^14^C]Gly, and the production of ^14^CO_2_ was measured over time. The data show that l-Ser and l-Thr were catabolized to CO_2_, but interestingly no Gly, under the conditions of the assay (Fig. 5 C). To track how much of the taken-up l-Ser, l-Thr or Gly offered to the cells at saturating concentration was oxidized to CO_2_, its production was quantified over time. After 3 h of incubation, 65.1% of the amount of l-Ser and 21% l-Thr that were transported into the cells were fully oxidized to ^14^CO_2_, while no CO_2_ derived from [U-^14^C]Gly was detected (Table 3). From these data it was concluded that l-Ser and l-Thr can be subjected to mitochondrial oxidation.

**Table 3.**
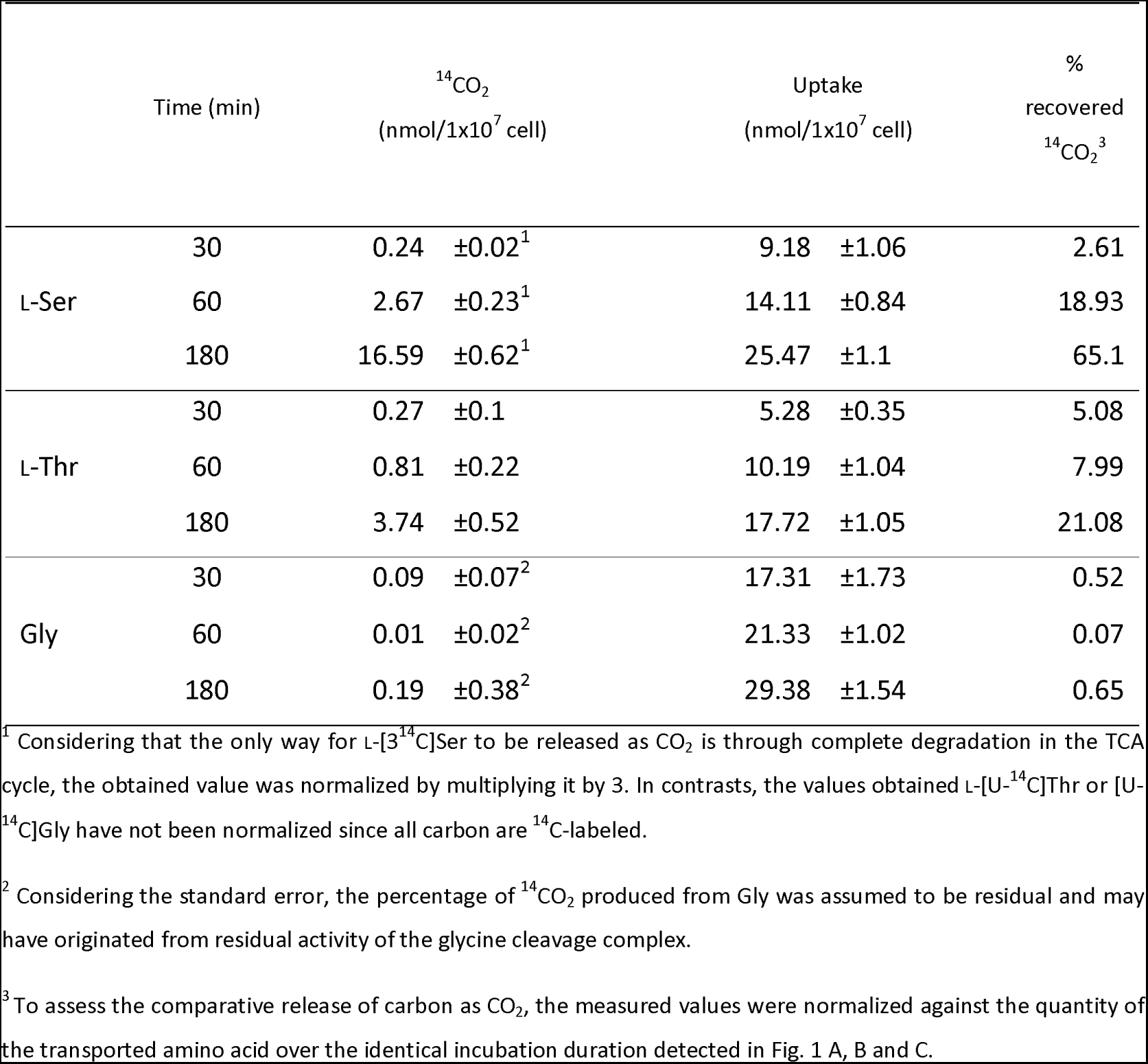
Time-dependent ^14^CO_2_ generation by Epi form from l-[3^14^C]Ser, l-[U^14^C]Thr and [U^14^C]Gly.

As the vast majority of mitochondrial oxidative processes are directed to ATP biosynthesis, it was investigated if this is also the case for l-Ser, l-Thr, and Gly. For this, Epi were submitted to NS by incubation in PBS for 30 h to deplete the intracellular ATP content, and their recovery by incubation in the presence of the substrates under investigation was assessed. The cells were recovered in PBS supplemented (or not, negative control) with 5 mM l-Ser, l-Thr, Gly or 5 mM His (control) for 3 h, and the intracellular ATP levels were determined. Parasites incubated with l-Ser and l-Thr exhibited a significant recovery of the intracellular ATP levels when compared to the negative control. This increase was comparable to those observed in parasites recovered with His. Interestingly, Gly could not restore intracellular ATP levels, which is in accordance with the absence of CO_2_ production (Fig. 5 D).

The observation that l-Ser and l-Thr can be taken up from the extracellular environment, support cell viability, produce CO_2,_ and support ATP biosynthesis, suggests that these amino acids are involved in oxidative phosphorylation (OxPhos). Thus, Epi subjected to NS (PBS) for 16 h, were transferred to MCR respiration buffer and recovered for 3 h by incubation in MCR supplemented (or not, negative control) with 5 mM l-Ser, l-Thr or His (positive control). The cells were then subjected to high-resolution oxygraphy, from which was derived the respiration parameters: **Routine respiration (R)**, rate of respiration that is not responsible for ATP production through OxPhos, **Leak respiration (L)**, the remaining O_2_ consumption measured when the ATP synthase is inhibited, **Electron-Transport Capacity (ETC)**, obtained after uncoupling the mitochondria by the addition of an uncoupler and **Residual Respiration Rate (ROX)**, obtained by the inhibition of complex III (Coenzyme Q-cytochrome C reductase), abolishing the flux of electrons through the respiratory chain (35). Our results demonstrate that after 3 h of incubation, l-Ser and l-Thr (but not Gly) were able to trigger O_2_ consumption (Fig. 6 A, B and C), when compared with the negative control (Fig. 6 E), and similarly to our positive control (Fig. 6 D). Furthermore, when **L** was subtracted from **R** to obtain the Free Routine Respiration (respiration rate directly related to the ATP biosynthesis by F_o_F_1_-ATP synthase) (36), l-Ser and l-Thr performed similarly as His (Fig. 6 G). Together, these results show the anaplerotic capacity of l-Ser and l-Thr, and thus, their ability to support ATP synthesis via OxPhos. Interestingly, and consistent with the CO_2_ and ATP levels measurements, Gly was not able to stimulate respiration in these cells.

**Figure 6.**
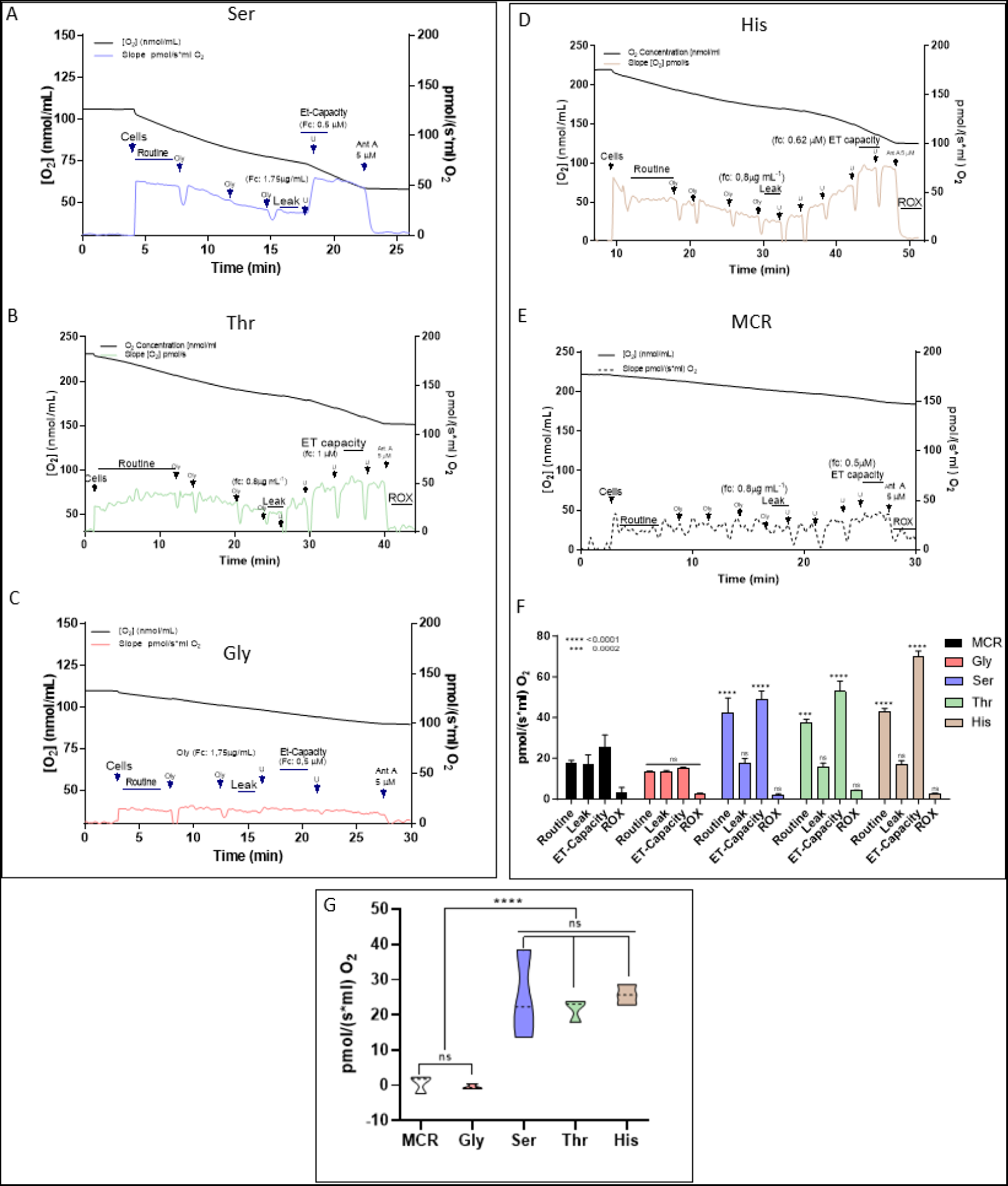
High-resolution respirometry in *T. cruzi* epimastigotes. Respiration rates were measured after 16 h of NS, when the parasites were recovered for 3 h in the presence of a substrate. A and B) respiration rates after 3 h of incubation with 5 mM l-Ser; C and D) respiration rates after 3 h of incubation with 5 mM l-Thr; E and F) respiration rates after 3 h of incubation with 5 mM Gly; G) positive control with 5 mM His; H). control without exogenous substrate; I) Free routine activity in epimastigotes recovered from NS with Gly, l-Ser and l-Thr. The Free Routine Activity (FRA) was obtained by subtracting the respiratory rates measured after addition of Oly. The graph is representative of this subtraction, using the average of the slopes obtained with the amino acids and after inhibition of F_o_-ATP synthase with Oly. The graphs are representative of three independent biological replicates. For graphs A to H, Two-way ANOVA with Tukey post-test was used for statistical analysis. **** P< 0.0001; *** P< 0.001. For graph I, one-way ANOVA with Tukey post-test was used. **** P< 0.0001.

Knowing that l-Ser stimulates both oxygen consumption and ATP biosynthesis, it is conceivable that this substrate could also contribute to establishing and maintaining the mitochondrial ΔΨ_m_ in *T. cruzi*. Using safranine O combined with controlled uncoupling of ΔΨ_m_ in the presence of valinomycin, the ΔΨ_m_ was quantified using the Nernst equation. For this experiment, the cells were permeabilized with digitonin in the presence of 2 mM l-Ser (Fig. 7 A). Given that l-Ser can form Pyr through Ser/ThrDH, the possible relationship between ΔΨ_m_ generated by l-Ser and the mitochondrial Pyr carrier (37) was tested by adding 5 µM UK-5099, a specific inhibitor of the Pyr mitochondrial transporter (MPC) (Fig. 7 A). As expected, l-Ser was able to build up and maintain ΔΨ_m_ in Epi. Consistent with the formation of Pyr through Ser/ThrDH, the generated ΔΨ_m_ was reversed in the presence of UK-5099. To rule out a possible off-target effect of UK-5099 altering the capacity of building up ΔΨ_m_, 2 mM Pro, a metabolite that can directly feed electrons into the respiratory chain (38,39), was added as a control (after UK 5099). As expected, the addition of Pro resulted in the reestablishment of the ΔΨ_m_. The quantification of the ΔΨ_m_ generated by l-Ser resulted in a ΔΨ_m_ of -190 mV, comparable to the positive control (l-Pro) and significantly different from the basal potential (∼ -90 mV) (Fig. 7 B). These results show that the conversion of l-Ser into Pyr is the main (or maybe the only) metabolic pathway linking this amino acid to ATP production through OxPhos.

**Figure 7.**
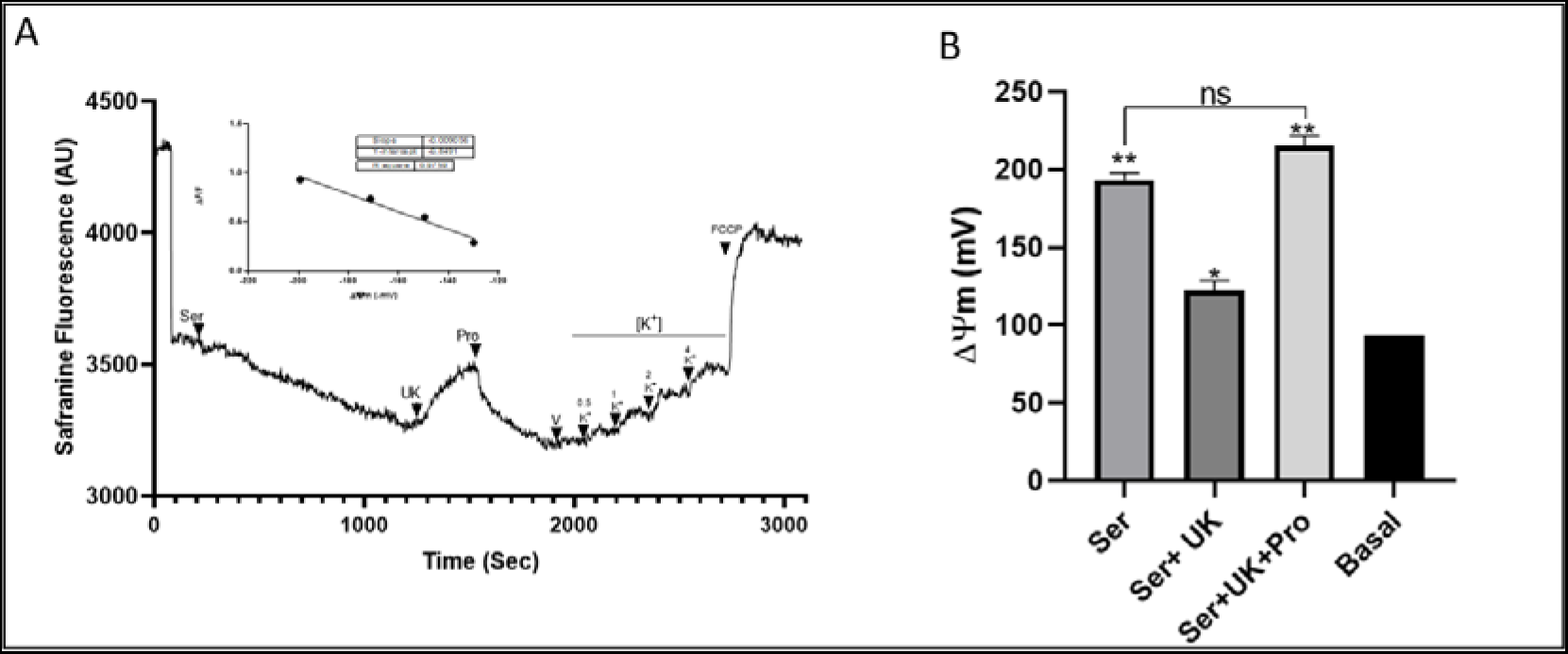
Quantification of ΔΨ_m_ in the presence of l-Ser and l-Pro. A) Representative trace of the ΔΨ_m_ quantification experiment by fluorometry, using safranine O in the presence of l-Ser, UK-5099 and Pro (positive control). Inset: calibration curve performed with K^+^ in the presence of 5 nM valinomycin. B) Quantification of ΔΨ_m_ under the previously mentioned conditions, using the Nernst equation (see materials and methods). The graphs are representative of three independent biological replicates. To compare the experimental conditions with the basal values and positive control (Pro), one-way ANOVA with Tukey’s post-test was used. ** P <0.001.

### Exometabolomics analysis to determine the excreted products from the metabolism of l-Ser

Given the bioenergetic relevance of the conversion of l-Ser into Pyr, the metabolic pathways involved in these processes were investigated by analyzing the products excreted by these cells when exposed to this amino acid. The exometabolome was investigated by using quantitative ^1^H-NMR analysis. For this, cells were incubated for 16h in the presence (or not, PBS) of 10 mM l-Ser. The spectra of proton resonances present in the samples were analyzed and quantified (Fig. 8 A and B).

**Figure 8.**
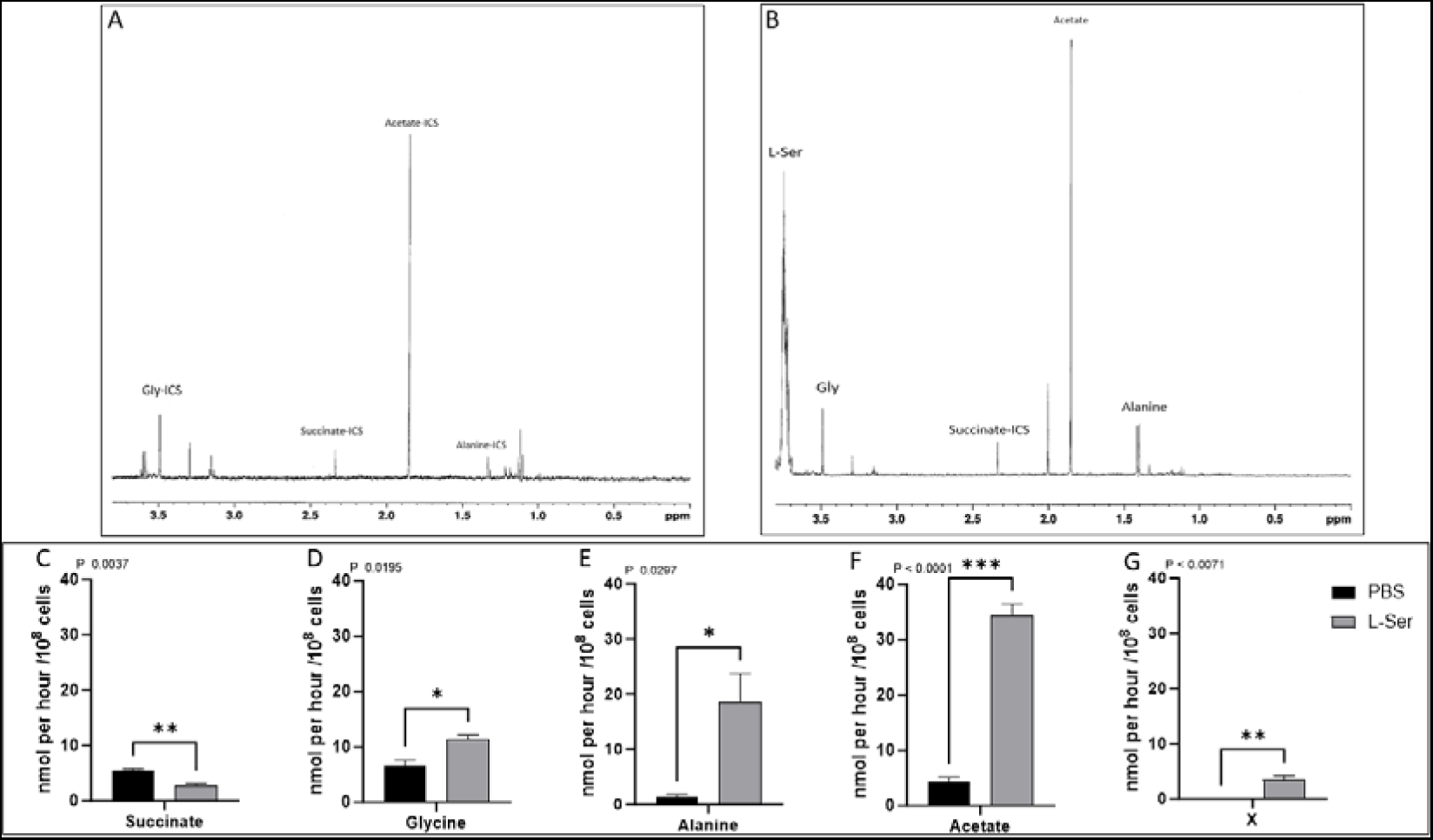
Excretion profile of metabolites from *T. cruzi* epimastigotes from l-Ser metabolism. (A) Proton resonance profile of metabolites excreted from parasites incubated for 16 h with PBS or (B) PBS supplemented with 10 mM l-Ser. (C, D, E, F and G) quantification of identified metabolites. ICS: Inner carbon sources. The graphs are representative of technical triplicates from three independent biological replicates. Two-tailed unpaired T test was used for statistical analysis. P <0.05 was considered statistically significant.

Our results show that the main excretion products from l-Ser metabolism are, in order of quantity, acetate, alanine and Gly, and a decreased excretion of succinate compared with excretion products of parasites without exogenous carbon source (Fig. 8). These data reinforce the idea that the metabolism of l-Ser in *T. cruzi* might occur through its conversion into Pyr, since both alanine and acetate are produced from Pyr (40). In line with this, an l-Ser/l-Thr dehydratase activity was detected in an Epi soluble extract (Fig. 9 A, B and C). To identify the molecular entity responsible for this activity, a search in the *T. cruzi* genome for an open reading frame encoding a putative l-Ser/l-Thr dehydratase was conducted. The identified gene (systematic number TcCLB.506825.70) was cloned and its product was heterologously expressed in *Escherichia coli* and purified (Fig. S2). The purified recombinant protein showed a l-Ser/l-Thr dehydratase activity (Fig. 9 D), confirming that the consumption of l-Ser for bioenergetics purposes in epimastigotes of *T. cruzi* can happen through its conversion into pyruvate.

**Figure 9.**
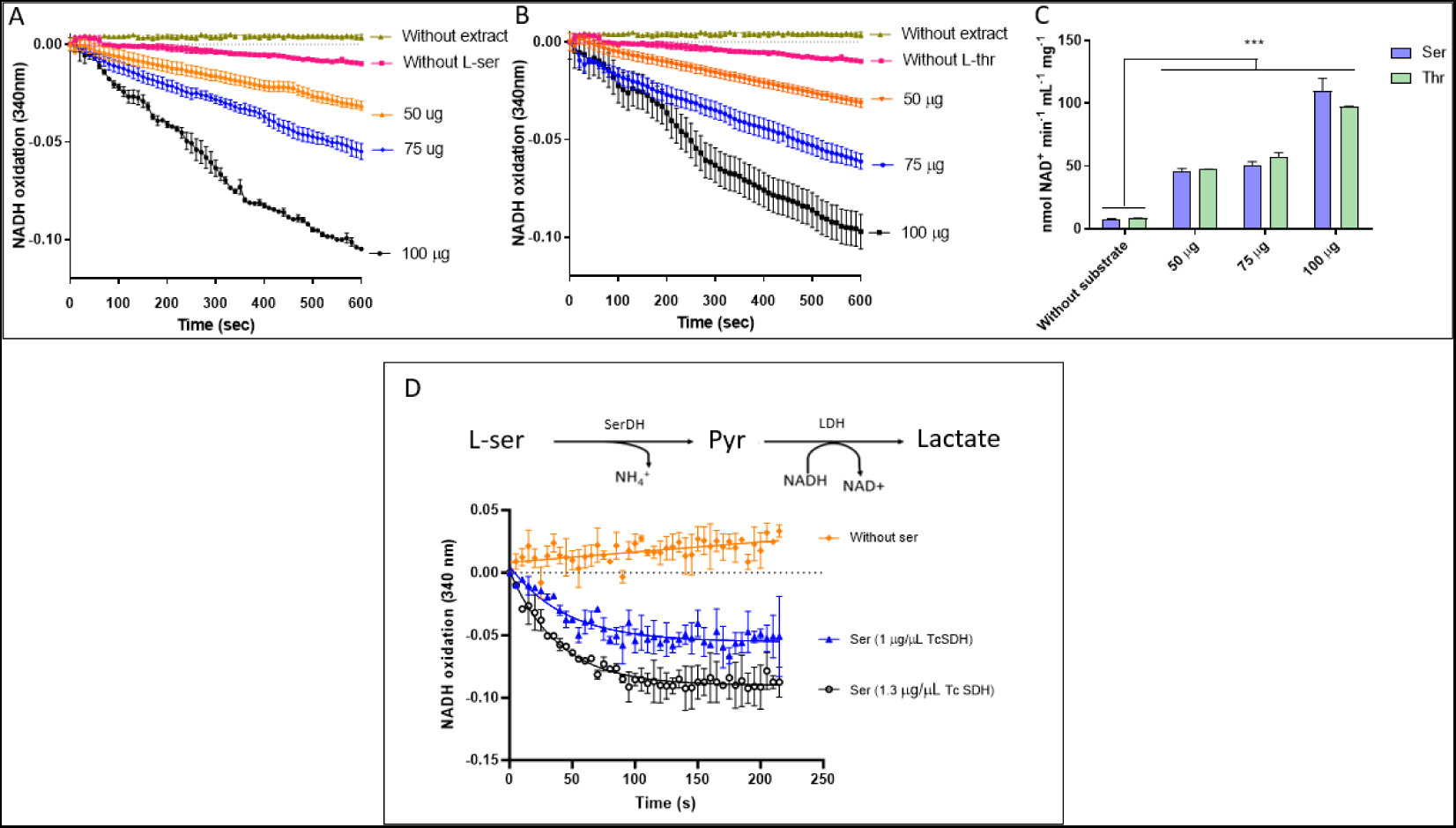
SerDH and ThrDH activities were measured by monitoring the decrease in absorbance at 340 nm (formation of NAD^+^) through coupling of their reactions with recombinant lactate dehydrogenase (LDH). Briefly, l-SerDH produces Pyr from l-ser, which through LDH produces lactate and oxidizes NADH. For ThrDH activity, 2-oxobutyrate is produced, which through LDH, produces 2-hydroxybutyrate also oxidizing NADH. (A) Activity using l-Ser as substrate: (B) Activity using l-Thr as substrate. C) NAD^+^ produced in the reaction at different concentrations of soluble parasite extract. (D) SerDH activity of the purified recombinant enzyme. The figure represents the average of 9 technical replicates in 3 independent experiments.

## DISCUSSION

*T. cruzi* presents an interesting case of metabolic adaptation since it can use a wide spectrum of compounds for its maintenance in vertebrate and invertebrate hosts. It has been demonstrated that after consuming preferentially Glc, it can switch to the use of amino acids, mostly proline (38,39,41), but also glutamate, aspartate, glutamine (42), histidine (32) or fatty acids (14,43,44). In the present work, we showed that it is also able to consume l-Ser and l-Thr as primary carbon and energy sources, while Gly can participate in its metabolism by providing carbons but not energy. In most organisms, l-Ser can be produced *de novo* from 3-phosphoglycerate (3PG), a glycolytic/gluconeogenic intermediate (45,46). However, *T. cruzi* lacks the last two steps of this pathway, suggesting that it relies on the uptake and/or a SHMT for obtaining l-Ser (21,47). SHMT can form l-Ser by its reverse reaction using Gly and 5,10-CH_2_-THF as substrates. Similarly, while plants, bacteria, and fungi can synthesize l-Thr *de novo* from aspartate (48,49), *T. cruzi* lacks the first three steps of this pathway. It has been proposed that *T. cruzi*, like *T. brucei*, might be capable of synthesizing l-Thr from homoserine and acyl-homoserine lactones, constituting a “by-pass” in the pathway at the homoserine kinase step (2.7.1.39) (18). Given these deficiencies in the biosynthesis pathways of l-Ser and l-Thr, *T. cruzi* relies on transport activities to obtain these molecules, which (among trypanosomatids) has to date only been demonstrated in *T. brucei* (for l-Thr) (17) and *Leishmania amazonensis* (for l-Ser) (50).

### The uptake of l-Ser, l-Thr and Gly occurs by a shared H^+^/ATP-driven transport system

The transport of metabolites can be considered the first step in metabolic pathways (15). Throughout its life cycle, *T. cruzi* is exposed to l-Ser, l-Thr, and Gly which are present in the insect vector’s excreta (51,52), as well as in mammalian cells and plasma (53–55). Many amino acid transport systems have already been biochemically characterized in this organism (Table S3). Noticeably, most of them have functional properties compatible with the AAAP family, which groups H^+^/amino acid transporters and auxin permeases (56,57). Due to its relevant relationship with l-Thr, and Gly, many aspects of their uptake and metabolism were analyzed together in this work.

The *K*_M_ values for l-Ser and l-Thr are in the same range of values reported for Cys (58), Asp (59), and the low-affinity Arg transporters (60). The *K*_M_ value measured for Gly resembles those reported for l-Glu (61), l-Ile (31), l-His (32) and high-affinity l-Pro (system A) transporters (62). The *V*_max_ values obtained for l-Ser and l-Thr are within the range of those for the BCAA, l-His, l-Gln and l-Pro low-affinity (system B) uptake systems (31,32,34,62). Additionally, the *V*_max_ value for Gly is amongst the three highest ones reported for *T. cruzi*, together with l-Ala (63) and l-Leu (31). This could be related to the use of Gly as an osmolyte. For this function, rapid changes in intracellular Gly concentration are needed as a response to hyper- or hypo-osmotic stress (64–67).

Given the fact that all three amino acids use the same transport system, the uptake of l-Ser was used as a proxy for all of them. Importantly, the transport activity for l-Ser increased exponentially at temperatures between 25 and 40 °C, a range to which the parasites are naturally exposed within both insect vectors (which do not control their temperature and thus their bodies follow environmental temperature variations) and vertebrate hosts. This suggests that the environmental temperature may be a natural modulator of l-Ser, l-Thr, and Gly uptake. From the response to temperature, the *E*_a_ was derived, which appeared to be the second lowest among those reported for amino acid transport systems in *T. cruzi*, after the low-affinity Arg transporter (60). Accordingly, from the *E*_a_ value, the equivalent of at least 1.6 molecules of ATP into ADP plus Pi per l-Ser molecule would be required for the latter to be transported into the cells. Finally, our data show that this uptake system is mediated by an active transporter as shown for most of the other amino acids in *T. cruzi*. With His transporter as the only exception (32), the primary mechanism driving these processes is a transmembrane H^+^ gradient established by a proton-pumping ATPase located in the plasma membrane (68). This contrasts with the l-Ser uptake in *Leishmania amazonensis*, which appears to be powered by ATP, independent of external ions such as K^+^ or Na^+^ and remains unaffected by any of the tested ionophores (50). Noteworthy, l-Ser uptake changed throughout the parasite’s life cycle indicating variations in the amounts of l-Ser and l-Thr *T. cruzi* encounters during its life cycle. In mammals, l-Ser and l-Thr in plasma range from 100 to 200 μM, values above the *K*_M_ measured for l-Ser and l-Thr transport (53–55). In triatomines, little is known about the abundance of these amino acids in different compartments of the digestive tract or in excreta. However, it can be assumed that Epi could find an environment rich in amino acids for two reasons: i) during the blood meal, the triatomine has high proteolytic activity, efficiently breaking down whole blood proteins (mostly hemoglobin) and thus releasing free amino acids that could be used by the parasite; and ii) during the bloodmeal, the triatomine usually releases amino acids into the intestinal lumen through the Malpighian tubules (51,52).

### l-Ser and l-Thr are fully catabolized to CO_2_ and stimulate OxPhos in epimastigotes

Differently to Gly, l-Ser and l-Thr were fully catabolized to CO_2_ and were able to sustain the intracellular ATP levels and respiration when present as the only carbon source. It is worth noting that the study of the participation of Gly was motivated by the possible existence of two pathways leading to the production of CO_2_: i) a SHMT activity which could interconvert Gly and 5,10-CH_2_-THF into l-Ser and THF (tetrahydrofolate) has been previously demonstrated in *T. cruzi* (20,69); and ii) the existence in the *T. cruzi* genome of putative genes encoding the Gly cleavage complex (GCC) (without demonstration of enzymatic functionality so far). The absence of CO_2_ production from Gly rules out both possibilities in our experimental conditions. Interestingly, the activity in *Leishmania major* promastigotes was inferred from indirect assays, in which the incorporation of Gly carbon into DNA was evidenced, demonstrating that Gly indeed participates in one-carbon metabolism (69). Concerning l-Ser, the same work showed that, when it was removed from the semi-defined culture medium (RPMI - 1% SFB) a critical decrease in growth occurred even in the presence of excess Gly, leading to the conclusion that SHMT would not compensate for l-Ser demand for optimal proliferation (69). These results are in line with the absence of classical *de novo* l-Ser biosynthesis pathways in trypanosomatids. In *T. brucei*, this complex may be essential since the parasite does not have SHMT and 10-formyl tetrahydrofolate synthetase annotated in the database of its genome, suggesting that it must rely exclusively on GCC for folate metabolism. Finally, l-Ser was able to build up ΔΨ_m_ in a way that was dependent on the previously characterized mitochondrial Pyr transporter (37,70), indicating that: i) in the cytosol l-Ser is converted into Pyr to be transported into the mitochondrion; or ii) it is transported into the mitochondrion as l-Ser by the Pyr carrier if this transporter has a broad specificity.

### The *Trypanosoma cruzi* epimastigotes have Ser and Thr dehydratase activities

In most eukaryotic cells the catabolism of l-Ser and l-Thr happens through dehydration followed by deamination. In the case of l-Ser, the products are Pyr and NH_4_^+^, and the reaction occurs in two steps: a cleavage of the hydroxyl group in a pyridoxal-5’-phosphate (PLP)-dependent manner, forming aminoacrylate, followed by its non-enzymatic hydrolysis to form Pyr and NH_4_^+^ (71,72). The dehydration/deamination of l-Thr occurs in three steps in which the dehydration reaction produces 2-aminobut-2-enoate in a PLP-dependent manner, followed by the non-enzymatic formation of 2-iminobutanoate and a deamination forming 2-oxobutanoate and NH_4_^+^ (29). Both, Ser/ThrDH activities were found in Epi soluble extracts. It is worth noting that l-Thr can serve as a substrate for two other putative enzymes predicted in the *T. cruzi* genome: l-Thr dehydrogenase (E.C. 1.1.1.10, TcCLB.507923.10) and a 2-amino-3-ketobutyrate coenzyme A ligase (E.C. 2.3.1.29, TcCLB.511899.10), which have been demonstrated as crucial for acetate production and fatty-acid biosynthesis in procyclics of *T. brucei* (19,73). It is worth mentioning that acetate is a main excreted product (together with alanine and succinate) of *T. cruzi* metabolism when grown in a rich medium (1,4,74,75). According to the results reported herein, the catabolism of l-Ser can contribute to the production of acetyl-CoA and then excrete acetate. Considering that i) the mitochondrial acetate:succinate CoA transferase/succinyl-CoA synthetase (ASCT/SCS) cycle is operative in *T. cruzi* Epi (unpublished results, under study by our group), and ii) *T. cruzi* lacks putative genes for an acetyl-CoA hydrolase (ACH, EC: 3.1.2.1), the production of acetate from l-Ser could be coupled to substrate-level production of intramitochondrial ATP in addition to OxPhos. Furthermore, the excretion of Gly by l-Ser fed parasites suggests that Gly is excreted as an osmotic control mechanism (65,67).

The ATP production fueled by l-Ser depends on its conversion into Pyr by a Ser/ThrDH. The incubation of permeabilized cells that maintained the integrity of the mitochondrial and glycosomal membranes with this metabolite enabled the formation of a ΔΨ_m_. This, together with the fact that this formation of ΔΨ_m_ is impaired when the MPC is inhibited can be explained by two different hypotheses: i) *Tc*MPC would transport l-Ser in addition to Pyr; or ii) the MPC inhibitor UK-5099 inhibits a mitochondrial carrier for l-Ser as well. It is important that other subcellular locations for the production of Pyr from l-Ser, such as glycosomes, cannot be ruled out in intact cells since a peptide of the Ser/ThrDH was reported in the proteomic profile of *T. cruzi* glycosomes and an analysis of the sequence revealed a PTS1 glycosomal-targeting sequence (76).

## CONCLUSION

In conclusion, l-Ser, l-Thr and Gly can be taken up by the cells and contribute to the cellular homeostasis maintenance during nutritional stress, providing conditions necessary for the resumption of epimastigote proliferation. l-Ser and l-Thr can be further oxidized with the production of CO_2_, triggering O_2_ consumption, contributing to the maintenance of the inner mitochondrial membrane potential and powering ATP production through OxPhos. With the evidence collected throughout this work, and in accordance with the data collected in the literature and genetic databases, we propose a model for the energy metabolism of l-Ser and l-Thr as shown in Figure 10.

**Figure 10.**
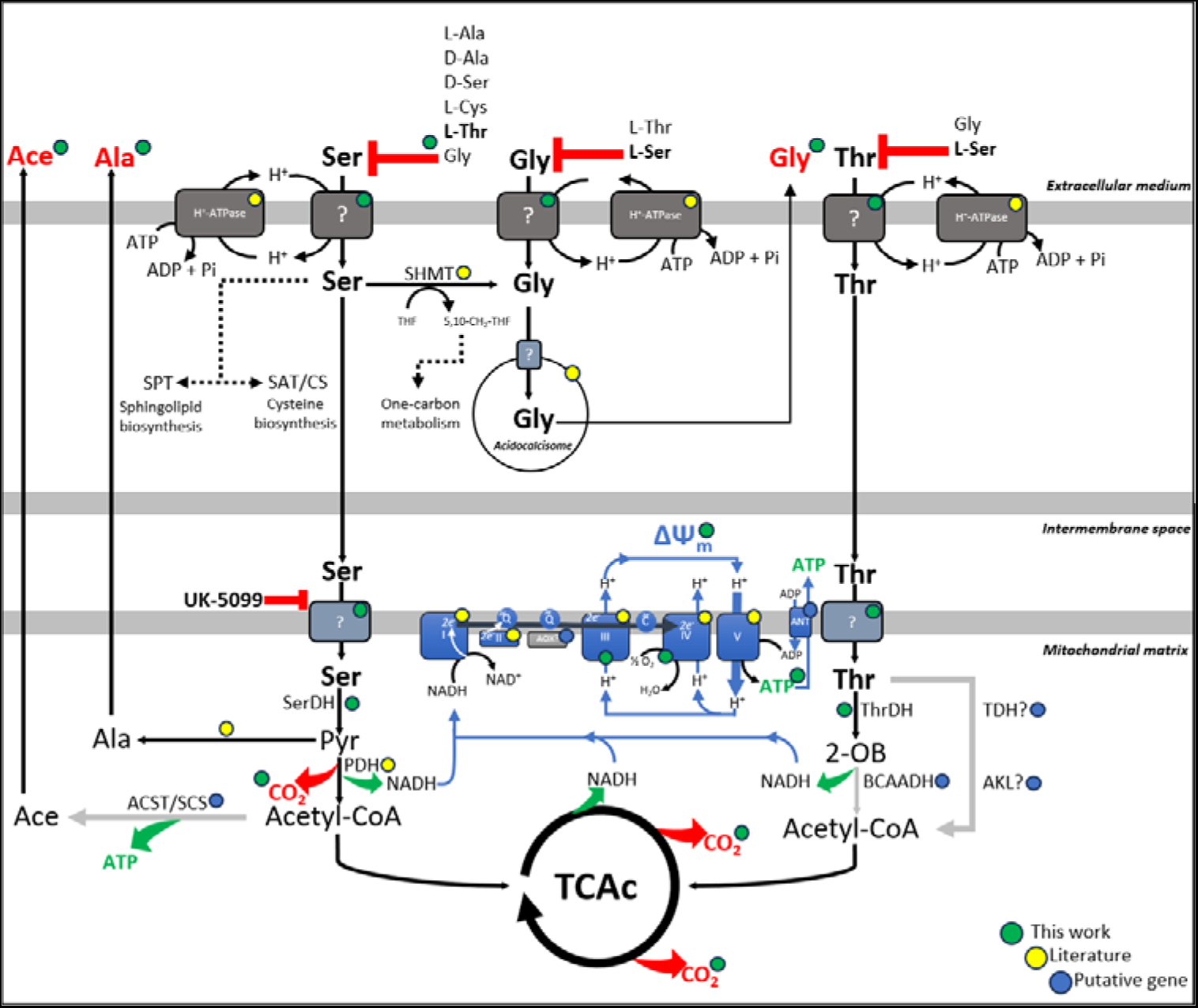
Metabolism of l-Ser and l-Thr in *Trypanosoma cruzi*. Metabolic steps are represented with different colors according to the origin of data: green-labeled steps correspond to data obtained in this work; yellow-labeled steps correspond to data obtained from the literature; blue-labeled steps correspond to inferred reactions according to annotations of the *T. cruzi* genome. Plasma-membrane H^+^-ATPase (68); Pyruvate Dehydrogenase (PDH) (77); mitochondrial electron-transfer complexes (CI-CIV) (35,78,79); alternative oxidase (AOX) (TcCLB.504147.180) It is important to mention that despite the identification of a putative gene for an AOX, the complex III inhibitor (antimycin A) abolish 99.9% of the respiration in *T. cruzi* Epi (35,78); F_1_F_o_-ATP synthase (35,78); adenine nucleotide translocase (ANT) (TcCLB.511249.10) (80); SPT (TcCLB.506405.50) (14,81); serine acetyltransferase and cysteine synthase (SAT/CS) (82,83); SHMT (21); Branched-chain alpha-keto acid dehydrogenase complex (BCAKDH) (Tc001047053506295.160; Tc001047053506853.50; Tc001047053507601.70; Tc001047053507757.70); l-Threonine 3-dehydrogenase (TDH) (TcCLB.507923.10); 2-amino-3-ketobutyrate coenzyme A ligase (AKL) (TcCLB.511899.10); acetate:succinate CoA transferase (ASCT) (TcCLB.504153.360); succinyl-CoA synthetase (SCS) (α: TcCLB.508479.340; β: TcCLB.507681.20).

## MATERIALS AND METHODS

### Reagents

l-[3-^14^C]Ser, l-[U-^14^C]Thr and [U-^14^C]Gly (0.1 mCi/ml) were purchased from American Radiolabeled Chemicals, Inc. (ARC [St. Louis, MO]). All other reagents were from Sigma (St. Louis, MO).

### Parasites

*T. cruzi* CL strain clone 14 epimastigotes were maintained in the exponential growth phase by subculturing them every 48 h in liver infusion tryptose (LIT) medium supplemented with 10% fetal calf serum (FCS) at 28 °C (84). For transport assays, exponentially growing parasites were washed three times with PBS (137 mM NaCl, 2.68 mM KCl, 8 mM Na_2_HPO_4_, and 1.47 mM KH_2_PO_4_, pH 7.2) and resuspended to a final density of 2 × 10^8^ cells/ml in PBS. To evaluate the ability of epimastigotes to use l-Ser, l-Thr and Gly in order to resist severe metabolic stress and to use it as an energy source, parasites in the exponential growth phase (5 × 10^7^ parasites per ml obtained from a 24h culture started at 2.5 × 10^7^ parasites/ml) were washed twice in 1 volume of PBS and incubated for 16h in 1 volume of the same buffer. After incubation, l-Ser, l-Thr or Gly was added to the cultures at a saturating concentration (5 mM) for its uptake, and different parameters of energy metabolism were determined, including cell viability, ATP production, oxygen consumption, and mitochondrial inner membrane potential.

### Transport assays

Transport assays were performed as described previously (31,32,34,62,63). Transport assays were initiated by the addition of 100 µl of 5 mM l-Ser, l-Thr or Gly in PBS to aliquots of parasites of 100 µl (2 × 10^7^ cells each, except when otherwise specified), traced with 0.4 µCi of radiolabeled amino acid. The uptake was measured at 28 °C for 1 min, except when otherwise specified. The transport reaction was stopped by addition of 800 µl of stop solution (50 mM amino acid in PBS, pH 7.4) prechilled at 4 °C, immediately followed by two washes with cold PBS. Background values in each experiment were measured by the simultaneous addition of each traced amino acid and stop solution.

### Competition assays

Competition assays were performed by measuring l-Ser, l-Thr or Gly uptake at a concentration equivalent to the *K*_M_ in the presence of 10 times the *K*_M_ of each other amino acid (29). Briefly, 100-µl aliquots of parasites containing 2 × 10^7^ cells were incubated with the transport solution supplemented with the presumably competing metabolite for 1 min. The results obtained were expressed as inhibition percentages in relation to a control (the same experiment without the competitor).

The effect of a H^+^-dependent plasma membrane potential on the l-Ser uptake was evaluated in parasites treated with 0.5 µM of the protonophore carbonyl cyanide *m*-chlorophenyl hydrazone (CCCP). As previously reported, protonophore treatment can affect an uptake process due to the disruption of the H^+^ gradient across cellular membranes (if the uptake is performed through a H^+^/metabolite symporter) or to the decrease of intracellular levels of ATP due to its rapid consumption by the mitochondrial F_1_F_o_-ATP synthase, which in a low-mitochondrial-membrane-potential situation hydrolyzes ATP to pump H^+^ to reestablish the mitochondrial inner membrane potential. To discriminate between both effects, a control was performed with the addition of 5 µg/mL oligomycin A to FCCP-treated cells, which allowed simultaneous disruption of H^+^ membrane gradients while blocking the F_1_F_o_-ATPase.

### Analysis of transport assay data

The disintegrations per minute (*dpm*) corresponding to transported radiolabeled amino acid for each experimental point (*dpmi*) were calculated as *dpmi = dpme − dpmb*, where *dpme* is the average *dpm* from triplicates after 1 min of incubation in the presence of radiolabeled amino acid and *dpmb* is the average *dpm* from the background samples.

The amount of amino acid taken up by the cells was calculated as *AAi = dpmi [AA] v dpmst^−1^ t^−1^*, where *AAi* is the transported amino acid, [*AA*] is the amino acid nanomolar concentration, *v* is the volume of radiolabeled amino acid solution in the experiment, *dpmst* is the total *dpm* measured for each added radiolabeled amino acids, and *t* is the time of incubation measured in minutes.

### Estimation of the number of transporters by using the Arrhenius equation

Thermodynamically, a transporter is an enzyme that catalyzes the reaction AA_e_ => AA_i_, where AA_e_ is the extracellular amino acid and AA_i_ is the intracellular amino acid. To estimate the number of transport systems, the turnover of active sites (*k*_cat_) must be measured. *k*_cat_ is calculated as: *k*_cat_ = *V*_max_/no. of transporters (85). Our hypothesis posits that, at saturated substrate concentration, the rate-limiting step of the system is dissociation. Consequently, the Arrhenius equation can be employed to estimate the number of transporter sites at a given temperature as follows:

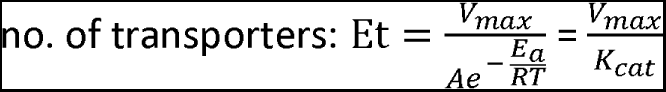

where *V*_max_ is the maximum rate achieved by the system at saturating substrate concentration at a given temperature, *Ae* is the pre-exponential factor, *E_a_* is the activation energy for the reaction, *R* is the universal gas constant, and *T* is the temperature of the reaction (in Kelvin).

### The transport of l-Ser and l-Thr in the life cycle forms of *T. cruzi*

After obtaining the different forms as described in references (33,34), the cells were resuspended in PBS and counted using a Neubauer chamber. The cell density was adjusted to a final density of 20 x 10^7^ cells/mL and distributed in aliquots of 100 µl (2 x 10^7^ cells each). The transport assay was initiated by adding 100 µl of 10 mM l-Ser or l-Thr labeled with l- [3-^14^C]Ser or l-[U-^14^C]Thr, respectively, prepared in PBS, pH 7.2. The initial velocity (*V*_0_) of the incorporation was measured at 28 °C (Epi and MTrypo) and 37 °C (Ama, Epi-like, and BTrypo) for 3 min. In all cases, the uptake of labelled amino acid was stopped by adding 800 µl of 50 mM of the corresponding unlabeled amino acid at 4 °C in PBS, and the parasites were washed by centrifugation at 10.000 x *g* for 2 min. The parasites were then resuspended in 100 µl of PBS and transferred to scintillation fluid. The samples were measured in a Perkin Elmer Tri-Carb 2910 TR scintillation detector to determine the *V*_0_ in each form.

### The contribution of l-Ser, l-Thr or Gly to recover cells subjected to a severe nutritional stress

To determine whether l-Ser, l-Thr or Gly can maintain the viability of epimastigotes of *T. cruzi* after a starvation period, the parasites were exponentially grown in LIT and stressed in PBS as described above. Briefly, the epimastigotes (5 × 10^7^ cells) were incubated for 24, 48, 72 and 96 h at 28°C in PBS plus 5 mM l-Ser, l-Thr or Gly to induce cell recovery. Separate treatments in histidine or PBS were used as controls. After recovery, the cells were washed in PBS and incubated with resazurin reagent to evaluate cell viability, as previously described (86).

Epi forms subjected to nutritional stress for 72h in PBS plus 5 mM l-Ser, l-Thr or Gly and incubated in PBS-HG (PPB + 5 mM l-His and 5 mM Glc, positive control) were transferred to LIT medium at a final density of 2.5 x 10^6^ parasites/mL, and then incubated 200 μL per well (experimental triplicates) in 96-well plates at 28 °C. The plates were read using an ELx800 Absorbance Reader® (Bio Tek®) where the dispersion of light by the culture was measured at λ 620 nm. Using a calibration curve with known parasite concentrations, assay values of light scattering by the culture (optical density, OD) were converted to number of parasites/mL.

### Quantification of ΔΨ_m_

Parasites in exponential growth phase (∼4-5x10^7^ parasites/mL) were centrifuged at 1,000 x *g* for 10 min at 4 °C and washed twice in ΔΨ_m_ buffer (250 mM sucrose, 10 mM HEPES, 250 µM EGTA, 2 mM NaH_2_PO_4_, 1 mM MgCl_2_ pH 7.2). Cells were adjusted to a density of 1x10^9^ parasites/mL. Aliquots of 50 µL were separated and permeabilized in the presence of 5 µM digitonin for 30 min. Safranin fluorescence was measured (EX WL: 495.0 nm, EM WL: 586.0 nm) in a cuvette fluorometer (F-7100 FL Spectrophotometer Hitachi High-Tech). Cells (5x10^7^ cells) were added to the cuvette in 2 mL ΔΨ_m_ buffer, 0.1% BSA (free of fatty acids), 12.5 µM safranin and 5 µM digitonin. The calibration curve was constructed under the same conditions as mentioned before. Then, 5 nM valinomycin was added, followed by titration using KCl. Finally, 1.25 µM FCCP was added to fully uncouple the membrane potential. The values were adjusted to the Nernst equation to estimate ΔΨm.

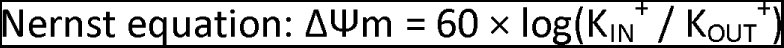

### Quantification of excreted metabolites by ^1^H-NMR analysis

Epimastigotes (1x10^8^/mL) were collected by centrifugation at 1.400 x *g* for 10 min, washed twice with PBS and incubated in 1 mL of PBS supplemented with 2 g/L NaHCO_3_ (pH 7.4). The cells were maintained for 6h at 27 °C in incubation buffer containing 5 mM l-Ser or no exogenous carbon sources. The integrity of the cells during the incubation was checked by microscopic observation. The supernatant (1 mL) was collected and 50 μl of maleate solution in D_2_O (10 mM) was added as an internal reference. ^1^H-NMR spectra were collected at 500.19 MHz on a Bruker Avance III 500 HD spectrometer equipped with a 5 mm Prodigy cryoprobe. The measurements were recorded at 25 °C. The acquisition conditions were as follows: 90° flip angle, 5.000 Hz spectral width, 32 K memory size, and 9.3 sec total recycling time. The measurements were performed with 64 scans for a total time of close to 10 min and 30 sec. The resonances of the obtained spectra were integrated, and the metabolite concentrations were calculated using the ERETIC2 NMR quantification Bruker program.

### ATP biosynthesis dependency of l-Ser, l-Thr and Gly

To evaluate ATP production with l-Ser, l-Thr or Gly as their sole energy source, the parasites (approximately 5 × 10^7^ cells/mL) were starved as described above and recovered or not (negative control) by incubation for 1 h in the presence of 5 mM His (as positive controls) or 5 mM l-Ser, l-Thr or Gly. The intracellular ATP concentration in each sample was determined after recovery by using a luciferase assay according to the manufacturer’s instructions (Sigma). ATP concentrations were estimated by using a calibration curve (ATP disodium salt, Sigma); luminescence (λ_570_ nm) was detected using a SpectraMax i3 plate reader (Molecular Devices, Sunnyvale, CA).

### CO_2_ production measurements

To measure the CO_2_ production from the TCA cycle during l-Ser, l-Thr or Gly catabolism, epimastigotes exponentially growing in LIT (5 × 10^7^ parasites/mL) were washed twice, resuspended in PBS and incubated in 5 mM l-Ser, l-Thr or Gly spiked with 0.1 µCi of radiolabeled amino acid for 0.5, 1, 2, 3 and 4h at 28 °C. To trap the produced CO_2_, pieces of Whatman filter embedded in 2 M KOH were placed on the top of the tubes in which the parasites were incubated. The filters were recovered and mixed with scintillation cocktail, and the K_2_^14^CO_3_ trapped on the paper was measured by using a scintillation counter. To assess the comparative release of carbon as CO_2_, the measured values were normalized against the quantity of the transported amino acid over an identical incubation period.

### Oxygen consumption

To evaluate respiration rates from the degradation of l-Ser, l-Thr and Gly, Epi (approximately 2.5 x 10^7^ cells/mL) were nutritionally stressed for 16h in PBS at 28 °C and recovered or not (negative control) for 3 h at 28 °C in the presence of 5 mM His (positive control) or 5 mM l-Ser, l-Thr and Gly as the only exogenous carbon sources. The parasites were added to the respiration buffer (MCR: 125 mM sucrose, 65 mM KCl, 10 mM HEPES NaOH, 1 mM MgCl_2_, 2 mM K_2_HPO_4_, pH 7.2). Subsequently, oligomycin A and FCCP were sequentially titrated. To verify residual respiration, a mitochondrial complex III inhibitor, antimycin A was added. Oxygen consumption rates were measured using intact cells in the high-resolution Oxygraph (OROBOROS, Oxygraph-2k, Innsbruck, AU). Data were recorded and treated with DataLab 8 software.

### Statistical analysis

Curve adjustments, regressions, and statistical analysis were performed with the GraphPad Prism 10 analysis tools. All assays were performed at least in biological triplicates, and the details of statistical analysis were added to each figure legend.

## Author contributions

MBA, RMBMG, MC, CGB, AMS, MB, FB: Conceptualization; Data curation; Formal analysis; Investigation; Methodology. MBA, AMS, FB: Validation; Visualization; Original draft; Review & Editing. AMS, FB: Funding acquisition; Project administration; Resources; Supervision.

---

**S3.** Overview of the biochemical characterization of amino acid transport systems to date in *Trypanosoma cruzi*.

| Amino acid | $K_M$ (mM) | $V_{max}$ (nmol·min<sup>-1</sup> per 20 × 10<sup>6</sup> cells) | $E_a$ (kJ/mol) | Ion and ATP dependence | | | | Ref. |
| --- | --- | --- | --- | --- | --- | --- | --- | --- |
|  |  |  |  | H<sup>+</sup> | K<sup>+</sup> | Na<sup>+</sup> | ATP |  |
| L-Alanine | 1.81 | 1.86 | 66.4 | yes | no | no | yes | (63) |
| L-Histidine | 0.12 | 0.24 | 55.25 | no | no | no | yes | (32) |
| L-Leucine | 1.07 | 0.94 |  |  |  |  |  |  |
| L-Isoleucine | 0.47 | 0.16 | 51.3 | yes | no | no | yes | (31) |
| L-Valine | 1.96 | 0.33 |  |  |  |  |  |  |
| L-Glutamine | 1.29 | 0.36 | 63.29 | yes | no | no | yes | (34) |
| L-Glutamate | 0.3 | 0.09834 | 52.38 | yes | no | no | yes | (61) |
| L-Arginine | 0.0485 | 0.03466 | 31.1 | ND | ND | ND | yes | (60, 88) |
| L-Aspartate | 0.032 | 0.0068 | ND | ND | ND | ND | ND | (59) |
| L-Cysteine | 0.0495 | 0.026 | ND | ND | ND | ND | ND | (58) |
| L-Proline A | 0.31 | 0.012 | 79.5 | yes | no | no | yes | (62) |
| L-Proline B | 1.36 | 0.31 | 18.3 e<br>176.5 | yes | no | no | yes |  |
| Glycine | 0.267 | 0.902 |  | yes | ND | ND | yes |  |
| L-Serine | 0.037 | 0.258 | 51.23 | yes | ND | ND | yes | This work |
| L-Threonine | 0.087 | 0.367 |  | yes | ND | ND | yes |  |

ND: no data

