## Supplemental material for "Elucidating the Transport Mechanisms and Metabolic Roles of Serine, Threonine, and Glycine in *Trypanosoma cruzi*"

**S1**

Computation:

$K_{cat} = Ae^{-\frac{E_a}{RT}}$

At 30°C, [S] at saturate concentration

R has the value of 8.3144621 x $10^{-3}$ kJ $mol^{-1}K^{-1}$

A = 1.34 x $10^{16}$ (from the Arrhenius Equation plot)

E = natural log base (2.718281828459)

$E_a$ = 51.23 kJ/mol

51,23 => kcat **1.995 x $10^7$ molecules $min^{-1}$ = 1.197 x $10^9$ molecules $s^{-1}$**

T = 303.15 K

$K_{cat} = Ae^{-\frac{E_a}{RT}}$

$K_{cat}$ Value obtained **1.995 x $10^7$ molecules $min^{-1}$ = 1.197 x $10^9$ molecules $s^{-1}$**

$V_{max}$ = 0.258 nmol/min per 20x$10^6$ cells

no. of transporters: $\mathrm{Et} = \frac{V_{max}}{Ae^{-\frac{E_a}{RT}}} = \frac{V_{max}}{K_{cat}}$

Et = $\frac{V_{max}}{K_{cat}}$ = 0.258/0.00198 = 0.00198 nmol per 20.$10^6$ cells

= 1.99 x$10^{-12}$ mol x 20.$10^6$ cells

= 9.9 x$10^{-13}$ mol x 10.$10^6$ cells

= 9.94x $10^{-5}$ amol x 10.$10^6$ cells

Average: 0.09938325 amol per cell

5.98 x $10^4$ active sites per cell

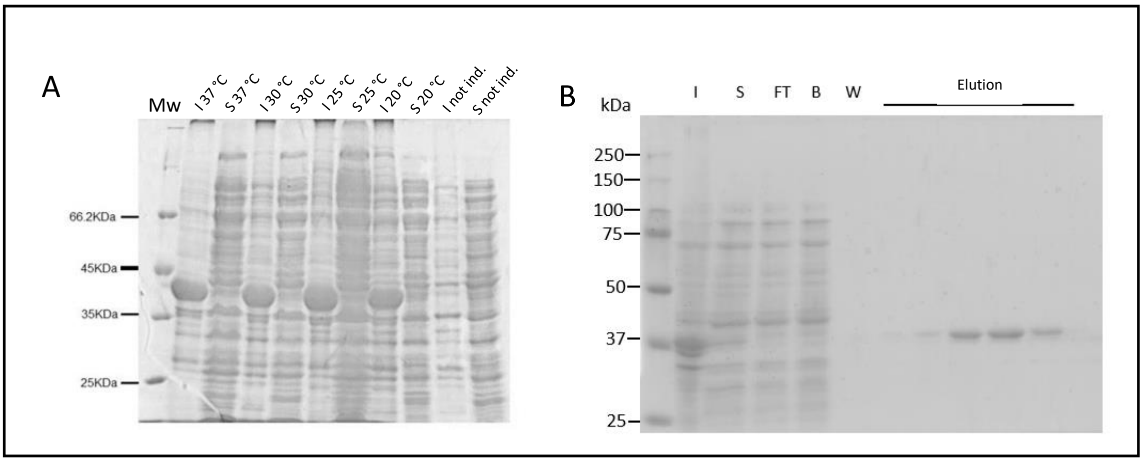


**S2.** (A) induction of the recombinant protein as a function of temperature. (B) SDS-PAGE of the recombinant enzyme purification from *E. coli* BL21 Codon Plus extracts by affinity chromatography on a $Ni^{2+}$-NTA column; I: insoluble; S: Soluble; FT: Flow Trough; B: Binding; W: Wash.

**S3. Overview of the biochemical characterization of amino acid transport systems to date in *Trypanosoma cruzi*.**

| | | | | Ion and ATP dependence | | | | |
|---|---|---|---|---|---|---|---|---|
| Amino acid | $K_M$ (mM) | $V_{max}$ (nmol·min$^{-1}$ per 20 × $10^6$ cells) | $E_a$ (kJ/mol) | $H^+$ | $K^+$ | $Na^+$ | ATP | Ref. |
| L-Alanine | 1.81 | 1.86 | 66.4 | yes | no | no | yes | (65) |
| L-Histidine | 0.12 | 0.24 | 55.25 | no | no | no | yes | (32) |
| L-Leucine | 1.07 | 0.94 | | | | | | |
| L-Isoleucine | 0.47 | 0.16 | 51.3 | yes | no | no | yes | (31) |
| L-Valine | 1.96 | 0.33 | | | | | | |
| L-Glutamine | 1.29 | 0.36 | 63.29 | yes | no | no | yes | (34) |
| L-Glutamate | 0.3 | 0.09834 | 52.38 | yes | no | no | yes | (63) |
| L-Arginine | 0.0485 | 0.03466 | 31.1 | ND | ND | ND | yes | (62,95) |
| LAspartate | 0.032 | 0.0068 | ND | ND | ND | ND | ND | (61) |
| L-Cysteine | 0.0495 | 0.026 | ND | ND | ND | ND | ND | (60) |
| L-Proline A | 0.31 | 0.012 | 79.5 | yes | no | no | yes | (64) |
| L-Proline B | 1.36 | 0.31 | 18.3 e 176.5 | yes | no | no | yes | |
| Glycine | 0.267 | 0.902 | | yes | ND | ND | yes | |
| L-Serine | 0.037 | 0.258 | 51.23 | yes | ND | ND | yes | This work |
| L-Threonine | 0.087 | 0.367 | | yes | ND | ND | yes | |
